# A quantitative model of sporadic axonal degeneration in the *Drosophila* visual system

**DOI:** 10.1101/2021.10.13.464240

**Authors:** Mélisande Richard, Karolína Doubková, Yohei Nitta, Hiroki Kawai, Atsushi Sugie, Gaia Tavosanis

**Affiliations:** Deutsches Zentrum für Neurodegenerative Erkrankungen e. V. (DZNE), 53127 Bonn, Germany; Department of Neuroscience of Disease, Brain Research Institute, Niigata University, Niigata 951-8585, Japan; LPIXEL Inc., 100-0004, Tokyo, Japan; LIMES Institute, University of Bonn, 53115 Bonn, Germany

## Abstract

In human neurodegenerative diseases, neurons undergo axonal degeneration months to years before they die. Here, we developed a system modelling early degenerative events in *Drosophila* adult photoreceptor cells. Thanks to the stereotypy of their axonal projections, this system delivers quantitative data on sporadic and progressive axonal degeneration of photoreceptor cells. Using this method, we show that exposure of adult flies to a constant light stimulation for several days overcomes the intrinsic resilience of R7 photoreceptors and leads to progressive axonal degeneration. This was not associated with apoptosis. We furthermore provide evidence that loss of synaptic integrity between R7 and a postsynaptic partner preceded axonal degeneration, thus recapitulating features of human neurodegenerative diseases. Finally, our experiments uncovered that neurotransmission to postsynaptic partners of R7 and their response are required to initiate degeneration, suggesting that postsynaptic cells signal back to the photoreceptor to maintain axonal structure. This model can be used to dissect cellular circuit mechanisms involved in the early events of axonal degeneration, allowing for a better understanding of how neurons cope with stress and lose their resilience capacities.

## Introduction

A remarkable property of neurons is their resilience. While most cells in the human body undergo frequent turnover, neurons in the central and peripheral nervous system can remain active and functional for decades (Sender and Milo, 2021). Nonetheless, in neurodegenerative conditions and during ageing, this resilience is overcome and progressive degeneration starts (Erkkinen et al., 2018; Hou et al., 2019). In this context, the initiation of neurodegeneration at the cellular level is represented by the shift from a resilient state to an unstable condition that ultimately leads to neuronal death. A small fraction of patients that develop commonly occurring neurodegenerative diseases, such as Alzheimer’s disease or Parkinson’s disease, display a familial predisposition (Brown, 1997; Schiesling et al., 2008; Tang and Gershon, 2003). Nonetheless, the vast majority of cases of neurodegenerative diseases do not have a clear familial history; they are thought to be sporadic, whereby ageing represents a major risk factor (Erkkinen et al., 2018; Hou et al., 2019). While developmental defects could underlie the onset of sporadic cases, a large body of evidence suggests that imbalanced neuronal activity leads to loss of neuronal resilience and triggers neurodegeneration (Arendt et al., 2017; Busche and Konnerth, 2016; Keogh et al., 2018; Palop and Mucke, 2016; Palop et al., 2007; Sosulina et al., 2021).

At the cellular level, the process of neuronal degeneration starts with the gradual loss of axons and dendrites (Adalbert and Coleman, 2013; Kweon et al., 2017). Axonal loss precedes degeneration of neuronal cell bodies by months to years, through a process of retrograde degeneration, termed ‘dying back pathology’ (Cavanagh, 1979; Neukomm and Freeman, 2014; Tagliaferro and Burke, 2016). Therefore, for potential intervention and to better understand the early phases of sporadic neurodegenerative phenomena, defining the initiation of axon damage is of great importance. Nonetheless, only recently these early phases have started to become more approachable for investigation in humans (Ahmad and Liu, 2020; Bondi et al., 2017; Bridi and Hirth, 2018; Grenier et al., 2020). Complementary work in model systems indicates that axonal ‘dying back’ in neurodegeneration shares some similarities with the active process of injured axon degeneration, which requires metabolic support and involves the interplay of Sarm, Axed and Wnk kinases (Coleman and Höke, 2020; Figley and DiAntonio, 2020; Gerdts et al., 2015; Izadifar et al., 2021; Neukomm et al., 2017; Walker et al., 2017; Yang et al., 2015; Yaron and Schuldiner, 2016). In addition, increased oxidative stress and dysfunctional autophagy are associated with axonal degeneration in many models of neurodegeneration (Hoffmann et al., 2019; Lee et al., 2021; Wang et al., 2018). Such dysfunctions target primarily the synapse, at which a slow accumulation of aggregation-prone proteins results in proteostasis imbalance and affects synaptic terminal composition, organization and function (Klaips et al., 2018; Lim and Yue, 2015; Olivero et al., 2018; Ross and Poirier, 2004; Wang et al., 2017). Indeed, synaptic dysfunction is now considered a major initial trait of many neurodegenerative diseases (Bellucci et al., 2016; Colom-Cadena et al., 2020; Compta and Revesz, 2021; Jack et al., 2013; Lepeta et al., 2016). Importantly, synapse loss precedes cognitive decline in many neurodegenerative diseases and is closely correlated with pathology progression (Colom-Cadena et al., 2020; Henstridge et al., 2018).

Invertebrate models, including *Drosophila* have greatly contributed to our understanding of neurodegenerative disorders, allowing for the identification and characterization of involved molecular factors (Bolus et al., 2020; Lu and Vogel, 2009; McGurk et al., 2015; Neukomm and Freeman, 2014; Şentürk and Bellen, 2018; Vanhauwaert and Verstreken, 2015; Wu and Lloyd, 2015). These models often rely on genetic manipulation of *Drosophila* orthologs of human neurodegenerative disease proteins or on transgenic expression of human proteins, linked to familial neurodegenerative diseases. However, a quantitative model for sporadic initiation of axonal degeneration is missing in this highly tractable model organism.

Here, we sought to develop a system in which axons of wild-type animals start to degenerate sporadically and reliably in defined conditions, with the aim of elucidating the mechanisms that trigger the switch from resilience to vulnerability of axons. For this purpose, we have developed a new fly model, combining important characteristics. First, we chose to work with photoreceptors, to allow for easy manipulation of activity by modulating the intensity or time exposure to light. Second, well-defined numbers of photoreceptors (PRCs) project their axon to higher visual processing centers. In particular, the R7 subset of PRCs projects to the medulla, where R7 axons distribute in highly ordered maps. The loss of individual axons is readily apparent in this checkerboard organization. Therefore, the initial cases of axon loss can be easily identified. Third, the circuit formed downstream of PRCs is described in detail at the EM level and sophisticated genetic tools allow the manipulation of individual relevant cell populations (Morante and Desplan, 2008; Takemura et al., 2008).

We show that exposure of flies to a constant light stimulation leads to a progressive light-dependent axonal degeneration in R7 PRCs, which was not accompanied by apoptosis. These axons slowly accumulated reversible damage up to a “point of no return”, which was reached at a different timepoint for individual axons and after which degeneration was initiated. Importantly, synaptic integrity between R7 and a medullar postsynaptic partner was lost before axons degenerated, thus recapitulating early features of human neurodegenerative diseases. Finally, neurotransmission to postsynaptic partners of R7 was required to initiate degeneration, suggesting that a postsynaptic signal elicited by prolonged neurotransmission triggers presynaptic axon degeneration.

## Material and methods

### Fly strains and light treatment

Flies were kept on standard medium in an incubator at 25 °C and 60% humidity (CLF Plant Climatics, Wertingen, Germany) and allowed to develop with a 12 hours of light/12 hours of darkness cycle (LD). After eclosion, flies were collected within 24 hours and kept in food vials at 25 °C and 60% humidity. For LD treatment, flies were illuminated with a 12 hours of light/12 hours of darkness cycle. We noticed that a light intensity up to 10,000lux in the LD cycle was not deleterious to axons (Fig. 2I, I’, J, K). For constant light (LL) conditions, we used a LED light source (LED panel, 6 Watt, warm white, Heitronic, Germany) and placed the flies in food vials at light intensity of 10,000lux, the intensity being measured at the top of the food in the vials and in the direction of the light source (photometer: Volt Craft MS-1300 Conrad Electronics, Germany). For constant darkness (DD), vials were wrapped in aluminium foil and kept in the same conditions as for LL.

All genotypes utilized are listed in Supplemental table 1. The following lines were obtained from the Bloomington stock center: *GMRwhiteRNAi* (#32067), *GMR-Gal4* (#1104), *UAS-tubGFP* (#7374), *UAS-p35* (#5072), *lexAop-spGFP1-10,UAS-CD4spGFP11* (#64315), *UAS-dcr2* (#24648), *tubGal80*^*ts*^ (#7018, #7019), *w* (#3605), *ort*^*5*^ (#29637), *UAS-CD8GFP* (#32186). The *40DUAS* line was used as a transgenic control and was obtained at VDRC (#60101). *Rh4-lexA* was described in (Berger-Müller et al., 2013). Following lines were generous gifts: *ortc2b-Gal4* from Dr. Chi-hon Lee (Gao et al., 2008), *UAS-shi*^*ts*^ from Prof. Michael Pankratz (Kitamoto, 2001), *UAS-dTrpA1*(II) from Prof. Tanimoto (Hamada et al., 2008). For *shi*^*ts*^ experiments, control and *UAS-shi*^*ts*^ expressing flies were raised in LD at 20 °C. After eclosion adults were shifted to LL at 29°C. For *TrpA1* experiments, flies were raised in LD at 20 °C and adults were kept at 29 °C (test) or 20 °C (control) in DD.

### Brain preparation

For medulla imaging, brains were prepared in PBS according to (Sugie et al., 2015, 2017) and fixed for 50 minutes in 4% paraformaldehyde. Brains were stained with mouse anti-Chaoptin (24B10, 1:25; Developmental Studies Hybridoma Bank) and incubated overnight at 4 °C in PBT containing 0.1% BSA. After washing, secondary Ab was incubated for 2 hours at room temperature (Alexa568-conjugated anti-mouse Ab, 1:400, Life Technologies). Brains were then mounted in Vectashield (Vector Labs) with the posterior side up after a second washing step. Insect pins of 0.1 mm diameter (Entomoravia, Czech Republic) were used as spacers to keep the brain in its original shape. Confocal microscopy was performed with a FV3000 (Figures 4E-O, 5, and 6 and Suppl. Fig. 4) or with a Zeiss LSM780 (Figures 1-3 and Suppl. Fig. 1-3). Images were processed using the IMARIS software (Bitplane) and the Fiji software (Schindelin et al., 2012).

### Axonal termini counts

In order to quantify the number of R7 axonal termini in medullas, confocal Z-stacks of ca. 100 μm in depth were acquired with a 40x objective (LSM780) or a 60x objective (FV3000). The Z-stack interval was fixed to 1µm, and following settings were used: 1AU, 1024 frame with LSM780 or 512 frame with FV3000, 8 bit with LSM780 or 16 bit with FV3000. We then reconstructed the stacks in 3D using the Imaris 9.7.2 software. Axonal termini of R7 PRCs project into the M6 medulla layer, which can be masked by using the Imaris “surface” function. Imaris allows for manual masking of axonal termini, which is highly time consuming. We thus created a python-based prediction software and trained it to generate a perfect mask of both tubulinGFP- and 24B10-positive R7 termini automatically. We trained the 2D-U-Net to generate a surface mask of three slices by inputting three slices as three channels, including the slice before and after the slice of interest. Then, during inference, only the mask of the centre slice was used. The training and testing datasets included images of both normal and affected medullas. A total of 16,114 images in 199 samples were used for training and 1,375 images in 16 samples were used for testing. By using three channels, the dice score improved from 0.815 to 0.847 compared to using a single channel. In order to use this software, a 3-step image pre-processing of the confocal stacks was needed in Fiji (Schindelin et al., 2012). We selected grey values in LUTs, the image type was changed into RGB colours and the image stack was further saved as an image sequence. After running our software, each folder was transformed into a corresponding folder with a predicted surface mask. Each folder was opened in Fiji as virtual stack and saved as a .tif file. In order to reconstruct the masked medulla M6 regions in 3D in Imaris, .tif stacks were converted into .ims files using the Imaris File Converter x64 9.7.2. For this, the .ims file corresponding to the masked M6 surface was opened in Imaris and transformed into 3D surface with the “surface” function (Scale: X:212, Y:212, Z:1, tick off smooth when creating surface). After that, the matching signal (tubulinGFP- or 24B10-positive axons) was added with the “add channel” function by selecting the .ims file equivalent to the initial folder and applying it on the created surface. In this step, the 3D-reconstructed full medulla is thus merged with the automatically generated surface. We then subtracted and visualised the M6 layer using the “mask channel” function (select in “channel tools” only the masked channel of interest). We subsequently aimed to count the number of individual axonal termini by using the “spot” function. To do so, we added four image processing steps into our protocol. We selected “gamma correction” values to 1.4, while “threshold cut off “ values were set up individually for each image as half of the automatically determined peak of intensity. In addition, we chose the “background subtraction” function with values suggested by the program and the “Gaussian filter” was selected as 0.6 µm. These processing steps allowed us to visualise axonal termini in better quality, thus facilitate the detection of termini by the “spot” function.

By using the “estimated XY diameter” function we estimated the size of a termini to 2µm (based on healthy axonal size in LD). Additionally, we aimed at avoiding the presence of off-targets, and thus used the “background subtraction” function, while keeping the quality thresholds manual: For images obtained with the LSM 780 confocal microscope, the quality threshold was determined as 2 for tubGFP signals and 1 for 24B10-positive axons; for images obtained with the Olympus confocal set up, the quality thresholds were adapted to 300 for 24B10 and 150 for the GFP signal. In order to provide highly accurate counts of axonal termini and reflect on early axonal loss which consists of up to only 5-10% of the total amount of axonal termini (Fig. 2J, K), we used the highly organised structure of medulla and took advantage of its axons organised in lines to manually evaluate the counts (Fig. 1H’-I’). Off-targets were then manually removed from the final count.

### Retinal staining/TUNEL assay

Eyes were removed from heads of *GMRwhite RNAi/w;GMR-gal4/+;UAS-tubGFP* flies in PBS and fixed in 4% paraformaldehyde overnight at 4°C. After washing in PBS Tween 0,1% (PBT), eyes were incubated in 50 mg/ml sodium borohydride in PBS for 20 minutes to remove pigments and subsequently washed in PBS and PBT. Cornea was removed and retinas were stained overnight at 4°C with Phalloidin-iFluor 488 (Biomol, 1/1000) in PBT BSA 0,1%. On the next day, retinas were washed with PBT and PBS and embedded directly in Fluoromount (Southern Biotech, Birmingham, AL) or further processed for TUNEL labelling. TUNEL staining was performed according to manufacturer’s instructions (In situ cell death detection kit TMR red, Roche). For the positive control, retinas were incubated with 300U/ml DNAse I in Tris pH7,5 1mg/ml BSA for 10 minutes at 25°C prior to TUNEL labelling. Retinas were washed in PBS and mounted with Fluoromount containing DAPI (Southern Biotech, Birmingham, AL).

### Eye pigment measurements

For brown (ommochrome) eye pigment measurements, we used a protocol adapted from (Mackenzie et al., 1999). 50 heads of 7-10 days-old flies were homogenized in 150 μl of 2 M HCl and 0.66% sodium metabisulfite (wt/vol). 200 μl of 1-butanol was added, and the mixture was placed on an orbital shaker at 150 rpm for 30 min before being centrifuged at 9000 *g* for 5 min. The organic layer was removed and washed with 150 μl of 0.66% sodium metabisulfite in dH2O and placed back on the orbital shaker for a further 30 min. The organic layer was removed and measured for absorbance at 492 nm. Absorbance was determined with Nanodrop One C spectrophotometer (Thermo Fischer).

Adult eye pictures were obtained with a Canon EOS 700D camera mounted on a Leica S8AP0 binocular and processed in Fiji (Schindelin et al., 2012).

### Circadian rhythm and sleep behavioral assay

The behavioral trials were carried out using the procedure described previously (Fogg et al., 2014; Yoshii et al., 2015), with some modifications. A DAM5 *Drosophila* activity monitor system (TriKinetics, Inc., Waltham, MA) was used to record locomotor activity in 1 min bins. Individual adult male flies of the genotype *GMR white RNAi/Y;GMRgal4;UAS-tubGFP*/+ from one to four days old were transferred to recording tubes containing 5% sucrose in 0.9% agar on one end. For constant light (LL) experiments, flies were entrained to a 12-h/12-h LD cycle (light: 4000 lux) at 25°C for 3 days. Subsequently, test flies were subjected to constant light (LL) conditions for 10 days and control flies to LD cycles for 10 days. Activity recordings were analysed using ActogramJ (Schmid et al., 2011). The sleep was defined as previously described (Huber et al., 2004).

### Statistical analysis

Statistical analyses were performed with GraphPad Prism 9.1.0 (GraphPad Software Inc.). All quantifications were performed by experimenters who were blind to the genotypes. The distribution of our data was determined using the D’Agostino & Pearson test and the Kolmogorov-Smirnov test (normality test was passed if P>0.05). For data following a Gaussian distribution, we used ordinary one-way ANOVA with Tukey’s multiple comparisons between groups. For experiments containing non-normally distributed data, we used the Kruskal-Wallis test and Dunn’s multiple comparisons between groups. For the sleep experiment, we used the Mann-Whitney U-test. P values above 0.05 were considered as non-significant. Statistical details are included in Supplemental table 2.

## Results

### Prolonged light exposure induces progressive axonal degeneration in photoreceptors

As a first step to establish a reliable setup for the induction of sporadic axonal degeneration in the nervous system, we studied the impact of prolonged light exposure on the visual system of adult flies. The *Drosophila* visual system consists of ∼750-800 independent unit eyes called ommatidia, which are organized in a crystalline-like arrangement (Tomlinson and Ready, 1987; Wolff and Ready, 1991). Each ommatidium is composed of eight PRCs that detect light and project retinotopically into the optic lobe, where visual processing occurs. Photoreceptor R1-R6 project axons to the first neuropil of the optic lobe, the lamina, while R7 and R8 contact their postsynaptic partners in the second neuropil, the medulla (Fischbach and Dittrich, 1989; Takemura et al., 2008). While cell death in the fly retina has been used extensively in neurodegeneration studies (Lenz et al., 2013), here we chose to monitor PRC axonal termini to concentrate on axonal degeneration. In particular, we focused on R7 PRCs, since their axon termini all reach a precise medulla layer (M6), where they display a highly ordered distribution and constant numbers (∼750-800) (Figure 1A) (Fischbach and Dittrich, 1989; Takemura et al., 2008). R7 axons and axonal termini can be readily visualized with antibodies against chaoptin (24B10) (Fujita et al., 1982) or by labelling PRCs genetically with *GMR-Gal4* driving expression of *UAS-tubulinGFP* (Fig. 1 B-E). Expression of *tubulinGFP* was detected along both R7 and R8 axons when imaging the medulla in a dorsal to ventral orientation (Fig. 1B, D) (Sugie et al., 2015). To obtain reliable counts of R7 axon terminals only, we first generated Z-scans of adult optic lobes, including the entire medulla, by imaging them in the posterior to anterior orientation (Fig. 1C). We then reconstructed all labelled axons in 3D and observed the R7 and R8 photoreceptor neurons project their axons side-by-side, until they terminate in distinct medulla layers (Fig. 1F). We then marked the R7 termini in M6 layer of the 3D reconstructed medullas to generate a 3D mask (Fig 1G), which we used to isolate the axon termini layer (Fig. 1H). This step is fully automated (see Materials and Methods). Each axon terminus was identified with a dot (see materials and methods) and the total number of dots was quantified (Fig. 1I). By imaging the medulla in this particular orientation and having extracted the M6 medulla region, we were able to reconstruct a complete checkerboard of R7 axonal termini, preserving their highly organized distribution. This allowed us to identify each missing axon (Fig. 1H’, I), particularly at the very early stages of neurodegeneration where only few axons are lost (Fig. 2). Thus, this technique allows for a precise and reliable counting of axonal termini of R7 in adult brains.

We exposed flies to a light intensity of ∼10,000 lux, an intensity corresponding to light measured in the shade of a summer day in the northern hemisphere (Schlichting et al., 2019). Photoreceptors implement multiple protective mechanisms to keep their activation level and their output to downstream neurons within a working range (Bai and Suzuki, 2020; Juusola and Hardie, 2001; Stavenga, 1995; Sugie et al., 2015, 2018). We found that R7 axons are highly resilient since R7 axon loss could be observed only starting after 3 weeks of continuous ∼10,000 lux light exposure (data not shown). This is primarily due to the fact that optical isolation of retinal ommatidia by the pigment cells protects retinal photoreceptors from excessive exposure to light (Bulgakova et al., 2010; Ferreiro et al., 2017; Lee and Montell, 2004; Schraermeyer and Dohms, 1993; Shoup, 1966). To thus facilitate the initiation of degeneration, we decreased pigment content in the retina by modulating the expression of the *white* gene, that codes for the pigment precursor transporter in pigment granules (Mackenzie et al., 2000; O’Hare et al., 1984; Pepling and Mount, 1990). In *white* mutant flies, degeneration started quickly and was widespread. To generate instead a situation in which axon degeneration is sporadic and progressive, we took advantage of a *GMR white RNAi* transgene to knock-down less efficiently *white* expression in the eye, thus yielding a yellow eye color (see Suppl. Fig. 1A, *white* progeny) (Lee and Carthew, 2003). In combination with *UAS-Dcr2*, this genotype displayed a homogenous eye color, independently of the inclusion of additional *w*^*+*^ transgenes (Suppl. Fig. 1).

**Figure 1.**
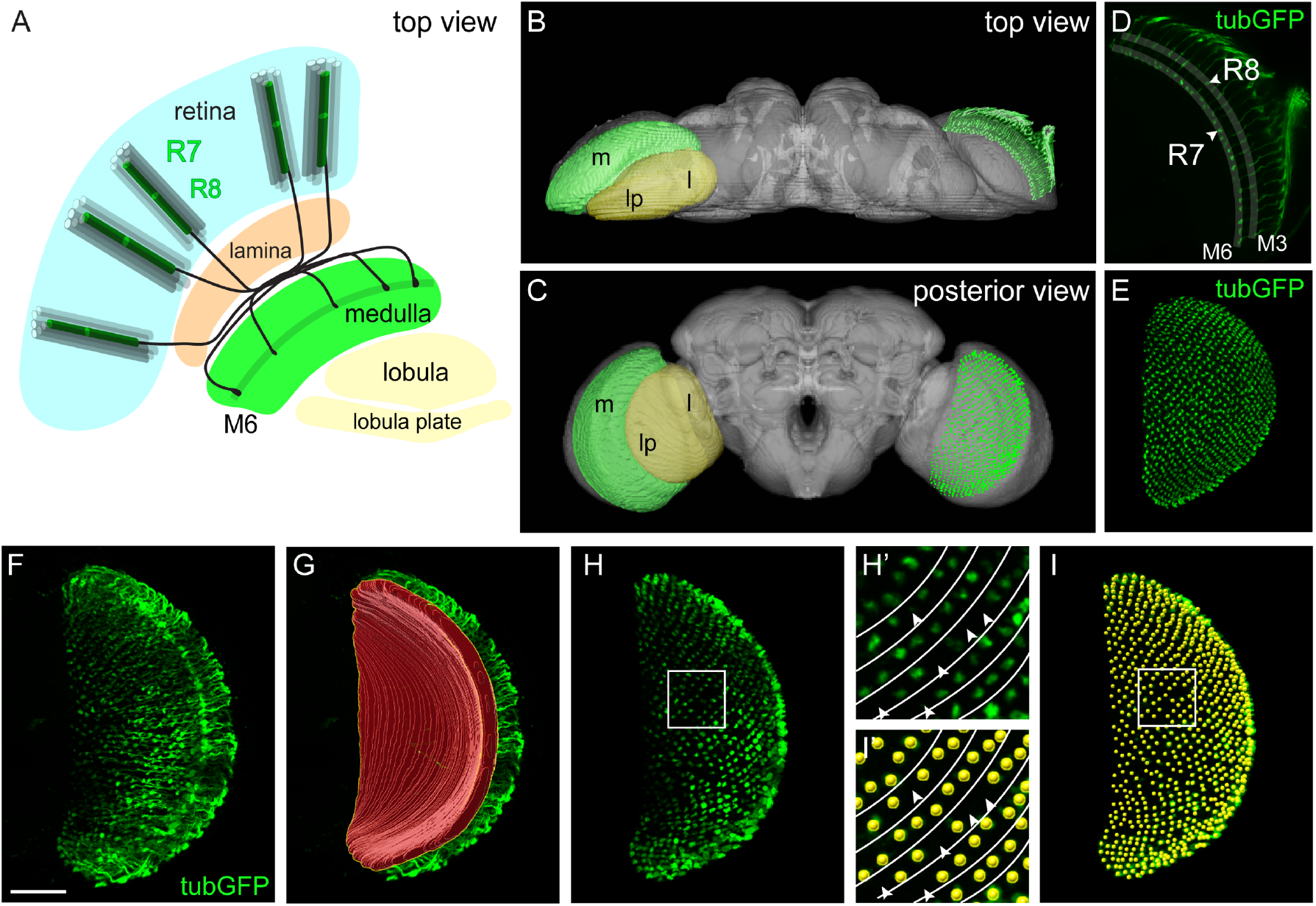
Quantitative analysis of sporadic axonal degeneration in *Drosophila* R7 photoreceptors. **A**. Scheme of the adult fly visual system. The retinal layer (blue) contains the cell bodies of the photoreceptor cells (PRCs), which are organized in ∼750-800 columnar units (ommatidia) all containing one PRC of each type (R1-R8). R1-R6 (gray columns) project their axons (not illustrated) to the lamina (orange). R7 and R8 PRCs (green) project their axons to the medulla (green); R7 termini (black) are found in medulla layer M6, while R8 terminate in M3 (not illustrated) (Fischbach and Dittrich, 1989). **B, C**. 3D image stacks of the adult fly brain (gray) depicting the positions of the medulla (green) and the lobula and lobula plate (yellow) in a top (B) or posterior (C) view (left part of the brain) as well as a Z-projection of R7 axons in a top view (B) or the R7 termini in the posterior view (C) (right part of the brain). **D**. Z-projection of tubulinGFP-labelled R7 and R8 axons imaged in a top brain view. R7 axons terminate in M6 (arrowhead), while R8 axons terminate in M3 (arrowhead). **E**. tubulinGFP-labelled R7 termini in a posterior brain view after extraction from a 3D brain reconstruction. The genotypes of this and all following Figures are included in Suppl. table 1. **F-H**. Identification of R7 axon termini in the M6 medulla layer. 3D-projection of tubulinGFP-labelled R7 and R8 axons in the medulla in a posterior brain view before extraction of R7 termini (F), 3D reconstruction of automatically created mask (red surface) to extract R7 termini in M6 (G) and R7 termini after extraction (H). **H’**. Close-up on R7 axonal termini in the medulla of flies expressing *UAS-tubulinGFP* driven by *GMR-gal4*. The organization of medullar R7 termini in rows (lines) facilitates the detection of missing axons upon prolonged light exposure (arrowheads). **I, I’**. A spot is assigned to each R7 axonal terminus (yellow). The total number of dots represents the amount of R7 axon termini in the medulla. Scale bar F-I: 50 μm.

In these flies, a small subset of R7 axonal termini started disappearing after 7 days of constant light exposure (LL) (Figure 2E, J, N). Axons degenerated progressively between 7 and 13 days of light exposure (Fig. 2J, F and G). By 13 days of light exposure, only ca. 60% of the 750-800 axonal termini found one day after eclosion (Fig. 2A, J) were still present (Fig. 2J, H, O). Axon loss occurred randomly throughout the neuropil, without obvious signs of regionalization. The numbers of axonal termini were unaffected by a cyclic exposure of 12 hours of light 10,000lux/12 hours of darkness (LD) for 13 days (Fig. 2I, J). To distinguish between loss of tubulin and loss of axons, we immunolabelled the medulla with antibodies against chaoptin (24B10), which decorates the membrane of R7 and R8 axons in the medulla, but only R7 in the M6 layer. 24B10-positive axons numbers progressively diminished upon LL exposure from day 7 to 13 in a way that mimics the loss of tubulin (Fig. 2K, E’-H’, N’, O’). They were unaffected by LD exposure (Fig. 2K, I’). Close-up optical sections at the axonal termini showed that after the first days in constant light, axonal termini firstly swelled (compare Fig. 2M and M’ with L and L’) before they started degenerating (Fig. 2N, N’). Since R7 axonal termini are organized in parallel rows, axon degeneration can be precisely and reliably monitored in this system (Fig. 2L-O’’). After 7 days in constant light, axon degeneration became apparent, with individual termini missing (Fig. 2N-N”). At 13dLL, axon degeneration was widespread in the medullas, and we observed both degenerating termini in which only chaoptin labeling was left (Fig. 2O-O”) and a large number of missing termini (Fig. 2O-O”).

**Figure 2.**
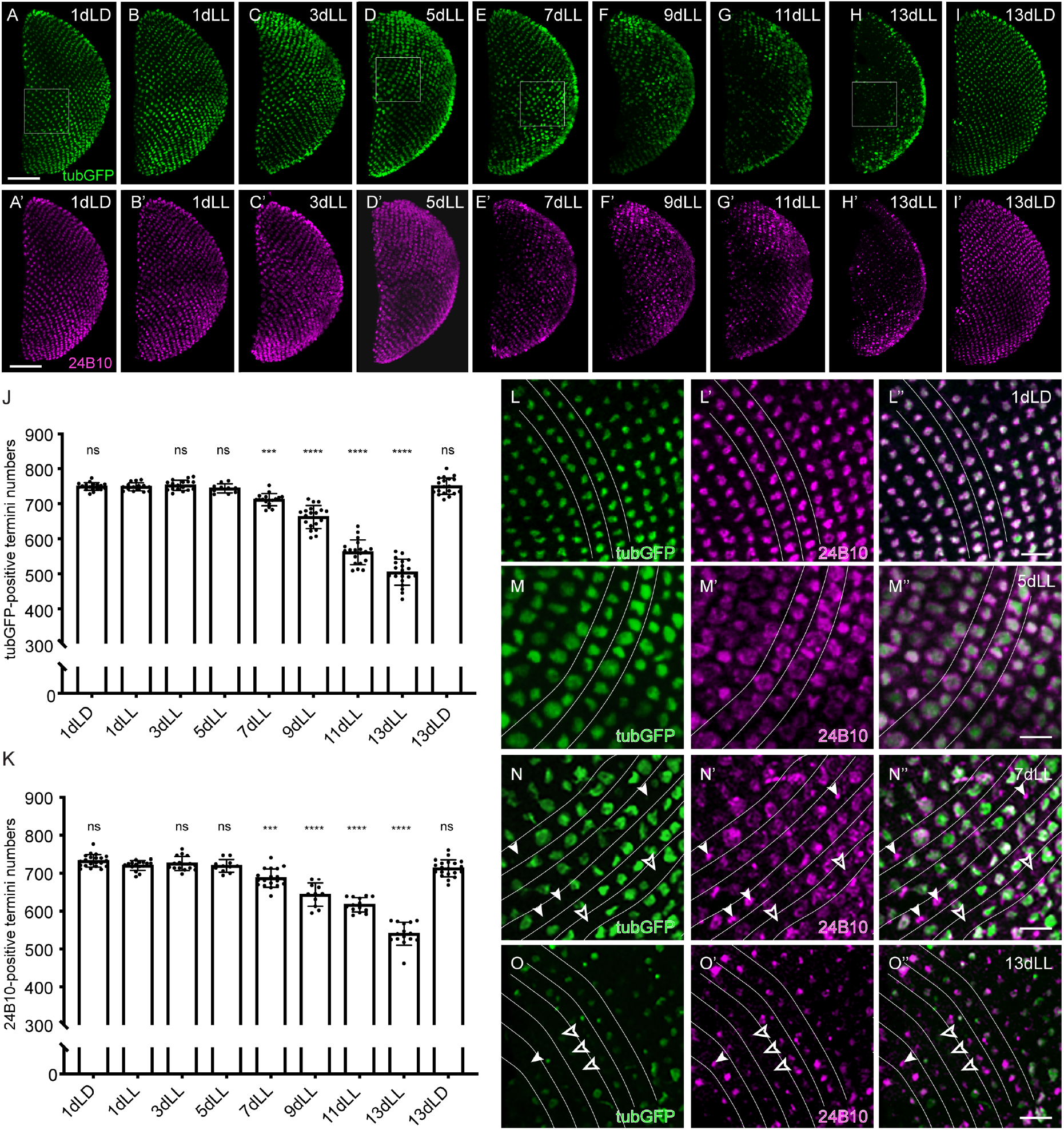
Light-dependent progressive degeneration of photoreceptor axons in the *Drosophila* medulla. **A-I and A’-I’**. Axonal termini of R7 in the medulla of flies exposed to constant light (LL) for various days or to control conditions for 1 day (A, A’) or 13 days (I, I’) in a 12 hours of light/12 hour of darkness (LD) cycle. (A-I) *UAS-tubulinGFP* driven by *GMR-gal4* (green). (A’-I’) Chaoptin (24B10) (magenta). Squares depict the position of the close-ups in L-O”. Scale bar: 50 μm. **J**. TubulinGFP-positive R7 termini counts in the medulla of flies exposed for various time to constant light (LL) or control conditions (LD). All statistics details for this and following graphs are included in Suppl. Table 2. **K**. 24B10-positive R7 termini counts in the medulla of flies exposed for various time to constant light (LL) or control conditions (LD). **L-O”**. Close-ups of R7 axonal termini in the medulla of flies at 1dLD (L-L”), 5dLL (M-M”), 7dLL (N-N”) and 13dLL (O-O”) labeled with *UAS-tubulinGFP* driven by *GMR-gal4* (green) (L-O); chaoptin (24B10) (magenta) (L’-O’); or merged (L”-O”). Filled arrowheads point to degenerating termini that have lost tubulinGFP expression but still retain the membrane marker 24B10. Empty arrowheads point at missing termini. Lines are drawn between rows of axonal termini. Scale bar: 10 μm.

Since constant light exposure abolishes circadian rhythm in the fruit fly (Konopka et al., 1989), we asked whether the light-dependent axonal degeneration observed in the medullas was due to a disruption of circadian rhythm. For this, we exposed flies to a weaker light intensity, that does not induce degeneration of R7 axons (4000lux, Suppl. Fig. 2A) and monitored their activity over a 10-day period either in LL (constant exposure of 4,000lux) or LD (12 hours of light at 4,000 lux/ 12 hours of darkness). As expected, flies in LL completely lost their rhythm after one day of constant mild light exposure (Suppl. Fig. 2B and C), showing that a mere disruption of cyclicity does not per se induce neurodegeneration in the medulla. In addition, we observed that this condition induced a slight increase of total sleep time in the LL flies, rather than sleep loss (Suppl. Fig.2 D).

Taken together, we developed a model of sporadic and progressive axonal degeneration in the medulla, in which it is possible to quantitively evaluate initial phases of axon loss.

### Reversibility of light-dependent axonal damage in R7 photoreceptors

We sought to identify the time point at which axonal degeneration was initiated. Five days of constant light exposure (5dLL) did not lead to R7 axonal loss in the medulla, but R7 axons underwent morphological changes, becoming enlarged (Fig. 2D-D’, J, K and M-M’’). We wondered whether axons at 5dLL are already damaged and whether a fraction of them is potentially committed to initiate degenerative processes. We thus placed those flies back into the dark (5dLL+6dDD) or exposed them to a light/dark cycle (5dLL+6dLD) and examined them at day 11 (Fig. 3A, E, F). Counts of tubulinGFP-positive termini among the medullas of 5dLL, 5dLL+6dDD or 5dLL+6dLD flies were not distinguishable from those of control animals (Fig. 3B, M). This is in strong contrast to the widespread axonal degeneration observed in 11dLL flies (Fig. 3C, M). Thus, the light-induced changes that had taken place in the R7 axons at 5dLL are reversible and the axons are not yet terminally committed to degenerate. Interestingly, the swelling presented by the axon termini in flies exposed to 5dLL was reversed in the 5dLL+6dDD condition, implying that axon swelling is a reversible modification at this stage (Fig. 3E). Axon swelling, however, was still present in the medulla of 5dLL+6dLD, indicating that prolonged D phases are needed for the swelling to disappear (Fig. 3F).

**Figure 3.**
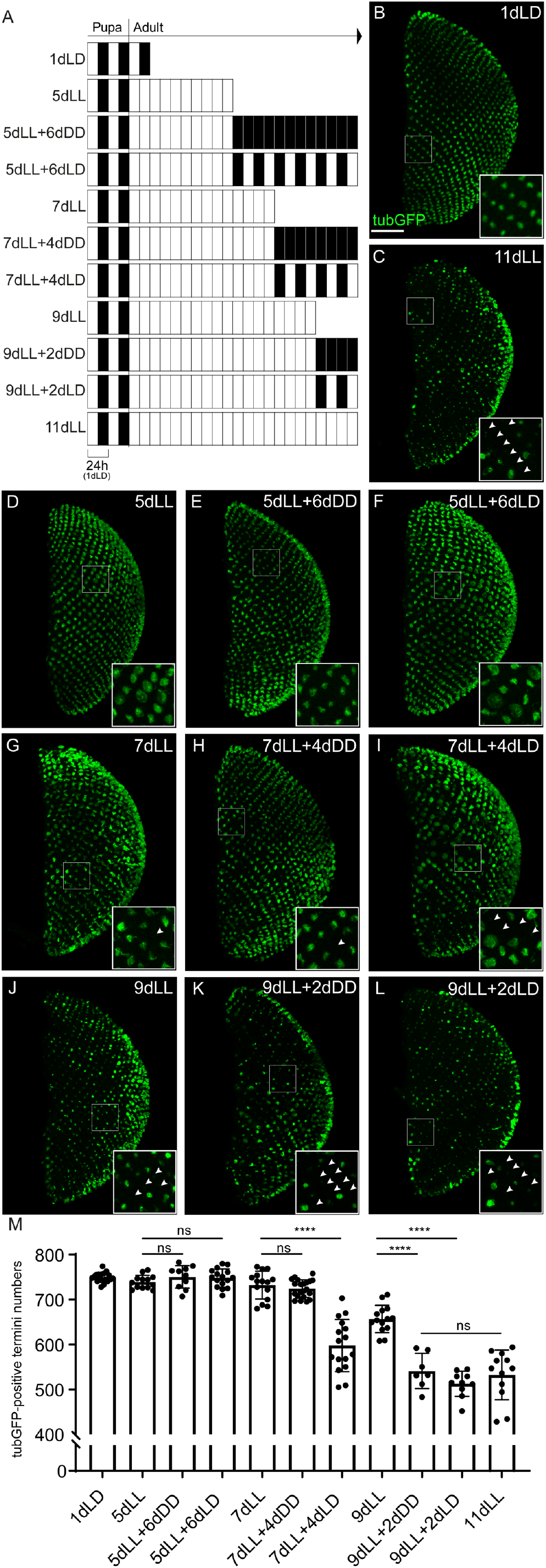
Reversibility of light-induced axonal damage in R7 photoreceptors. **A**. Scheme of the various illumination conditions used in this experiment. Development proceeded in a 12 hours of light (white)/12 hours of darkness (black) condition. At eclosion, adult flies were exposed to constant light for various time periods (white). A fraction of flies was put back to complete darkness (black) or in a light/dark cycle (black/white). **B-L**. Axonal termini of R7 in the medulla of flies expressing *UAS-tubulinGFP* (green) driven by *GMR-gal4* exposed to various light and dark cycles (A). We considered 1dLD as a non-degenerated condition (B), while 11dLL was chosen as the condition of strongest axonal loss (C). The position of the close-ups is depicted as a square and arrowheads point to missing axon termini. Scale bar: 50 µm. **M**. TubulinGFP-positive R7 termini counts in the medulla of flies exposed to various light/dark cycles.

At day 7 of continuous light exposure (7dLL), first individual axons started to lose the sensitive tubulinGFP signal (Fig. 3G). This situation was not worsened 4 days later if the flies were kept in the dark (7dLL+ 4dDD), suggesting that no additional axons had been terminally committed to degenerate at this stage (Fig. 3H). Axon swelling was also reduced in this condition. Nonetheless, if the 7dLL flies were placed in a light/dark cycle, progressive axonal degeneration became apparent (Fig. 3I, M). The final outcome was intermediate between that of 9dLL exposure and 11dLL exposure (Fig. 3M). This observation suggests that at 7dLL some axons have accumulated reversible damage and that further light exposure is necessary to drive these axons into degeneration.

After 9 days of continuous light exposure (9dLL), the medulla was obviously damaged, with many missing axons (Fig. 3J, M). Putting the flies back in the dark for 2 days (9dLL+ 2dDD) failed to slow down the progressive axonal degeneration and medullas of these flies were undistinguishable from those of flies exposed to 11dLL (Fig. 3C, K, M) or to those of flies exposed for 9dLL+ 2dLD (Fig 3L, M). Thus, at day 9 of light exposure, axon damage may be irreversible and a fraction of axons might be already committed to degeneration.

Taken together, these data suggest that R7 axons slowly accumulate reversible damage as a result of constant activation. Axons reach individually a level at which the accumulated damage initiates degeneration, a “point of no return”.

### Apoptosis is not associated with axonal degeneration

Axonal degeneration is an early event in NDs and precedes cell death (Adalbert and Coleman, 2013; Cavanagh, 1979; Neukomm and Freeman, 2014; Tagliaferro and Burke, 2016). To monitor whether axonal degeneration of R7 photoreceptor cells was a consequence of cell death or was preceding it in our model system, we firstly monitored the cornea. Eyes become rough in conditions in which individual PRCs are missing or when the number, arrangement, or identity of PRCs is modified (Basler et al., 1991; Tomlinson et al., 1987, 1988; Van Vactor et al., 1991). Flies exposed to 13 days of constant light (13dLL) did not show any detectable defects in corneal morphology, indicating that widespread cell death was not taking place at this stage (Fig. 4A). To address specifically whether individual PRCs might be undergoing apoptosis, we exposed flies to LD or LL treatments for various periods of time and performed TUNEL staining to detected apoptotic nuclei within the retinas (Fig. 4B-D and Suppl. Fig. 3A-I). Even after 13 days in constant light, we did not detect TUNEL-positive PRC nuclei (Fig. 4C), suggesting that apoptotic cell death had not started at a time point at which axonal degeneration was at full pace (Fig. 2H, H’, J) (see also Nitta et al., unpublished). Nuclear morphology was similar between controls and light-exposed flies (Fig. 4B, C). We observed a reported remodeling of the actin-rich rhabdomeres starting upon 5 days of light exposure (Suppl. Fig. 3F, G, H). Interestingly, R7 rhabdomeres seemed to be less affected than those of R1-R6 by this light-dependent actin remodeling after 5 or 7 days in constant light (Suppl. Fig. 3K, N). This type of actin redistribution in photoreceptors was already observed after short-term light exposure, and shown to be completely reversible in darkness (Kosloff et al., 2003). We thus tested whether we could reverse this phenotype by putting light-exposed flies (5LL or 7LL) back to darkness. A large majority of rhabdomeres recovered after 2 days in complete darkness (Suppl. Fig. 3L, O), thus suggesting that these changes at the rhabdomeres are not indicative of a permanent cell damage.

**Figure 4.**
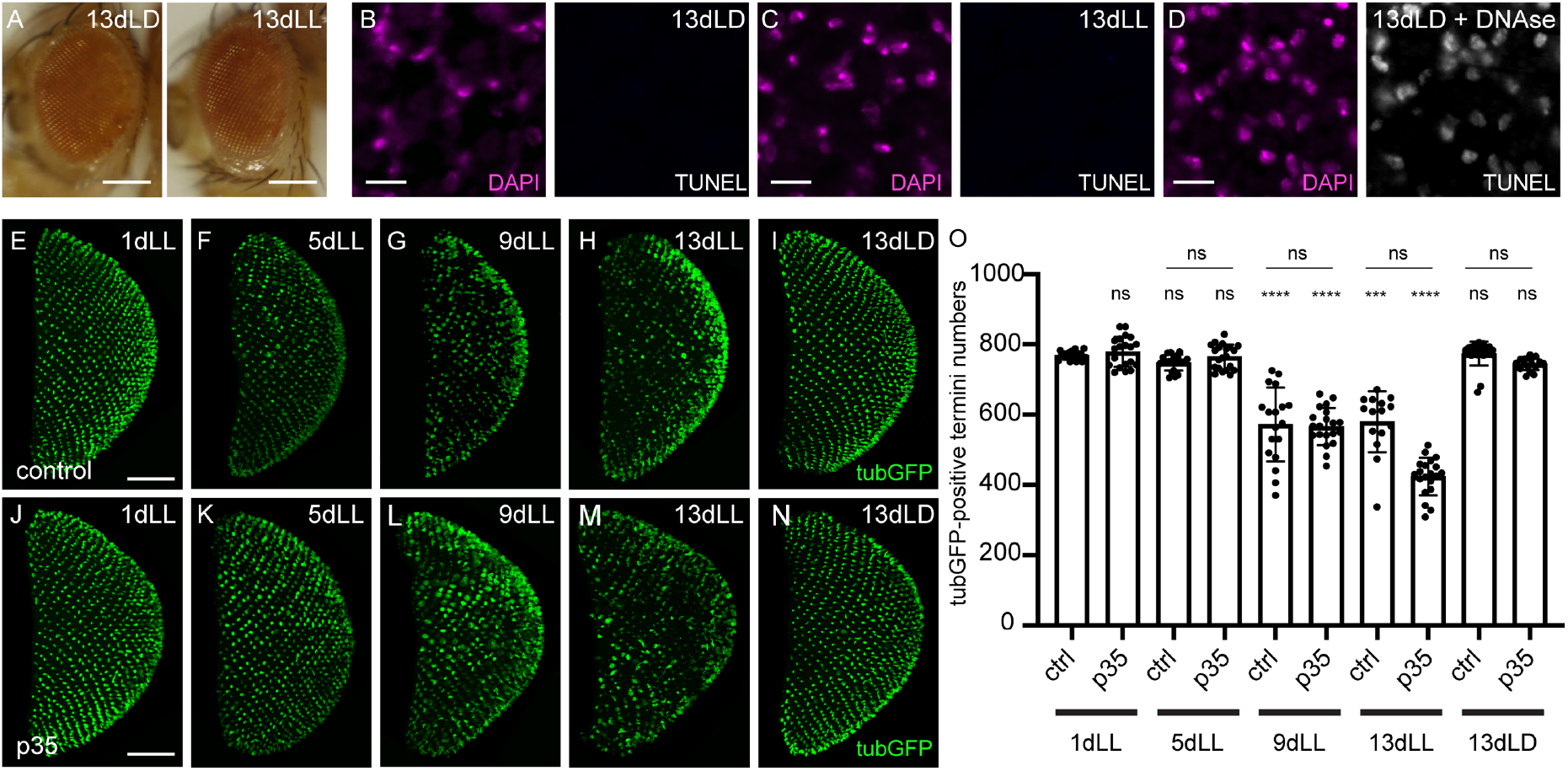
Apoptosis is not associated to axonal degeneration. **A**. Eyes of flies exposed for 13 days in constant light (LL) or to 13 days in a cycle of 12 hours of light:12 hours of darkness (LD). Scale bar: 250 μm. **B-D**. Optical cross-sections through the retina of adult flies exposed to 13 days of either control (13dLD in B) or constant (13dLL in C) light conditions. Eyes were stained with DAPI to mark the nuclei (magenta). TUNEL labelling (white) showed no signal except in the positive control, in which 13dLD eyes were submitted to a DNAse treatment before TUNEL detection (D). Scale bar: 10 μm. **E-N**. Axonal termini of R7 in the medulla of flies expressing p35 in J-N and control flies in E-I exposed for various time to constant light (LL) or to control conditions for 13 days in a 12 hours of light/12 hour of darkness (LD) condition. Green represents termini expressing *UAS-tubulinGFP* driven by *GMR-gal4*. Scale bar: 50 μm. **O**. TubulinGFP-positive R7 termini in the medulla of flies expressing p35 or of control flies exposed for various times to constant light (LL) or to control conditions for 13 days in a 12 hours of light/12 hour of darkness (LD) condition.

To independently validate that apoptosis was not involved in light-induced axonal degeneration during the time frame of our experiments, we expressed in PRCs the baculovirus caspase inhibitor p35 that suppresses apoptosis and analyzed axonal termini numbers after 1, 5, 9 and 13 days in LL (Fig. 4E-N) (Zhou et al., 1997). Expression of p35 did not modify termini numbers after 13 days in a light/dark control condition (Fig. 4I, N, O). Importantly, it did not rescue axonal degeneration at 9 or 13 days in constant light (Fig. 4L, M, O), indicating that apoptosis is not associated to the initial steps of axonal degeneration.

### Neurotransmission is involved in axonal degeneration

Recent evidence indicates that synaptic failure in AD is at least partly induced by neuronal hyperactivity in the early stages of the disease, and this mechanism could also be involved in developing a late-onset autosomal dominant inherited form of PD (Hector and Brouillette, 2020; Lucumi Moreno et al., 2021; Nuriel et al., 2017). We thus tested whether activity of PRCs is linked to the axonal degeneration described in our model. First, we tested if we could induce neurodegeneration in complete darkness by acutely activating R7 using the heat-sensitive *Drosophila* TrpA1 channel (Hamada et al., 2008; Pulver et al., 2009). Indeed, activation of photoreceptors by TrpA1 at 29ºC was sufficient to induce axonal degeneration in the dark (Fig. 5B), starting at 11 days after shifting to the restrictive temperature (Fig. 5C). Thus, simple activation of PRCs can induce axonal degeneration. As a next step, we expressed in R7 the temperature-sensitive transgene *UAS-shibire*^ts^ (*UAS-shi*^ts^), which prevents endocytosis at temperatures exceeding 29°C, thus rapidly stopping synaptic vesicle recycling and neurotransmitter release (Kitamoto, 2001). To avoid developmental defects caused by the leaky expression of *UAS-shi*^*ts*^, we included a *tubGal80*^*ts*^ transgene in the experimental genotype (McGuire et al., 2003). Flies thus developed at the permissive temperature and were shifted to the restrictive temperature at eclosion, allowing for *UAS-shi*^*ts*^ expression and dominant negative activity only at adult stages (see Material and Methods). In contrast to the degeneration observed in control flies (Fig. 5D, F), flies expressing *UAS-shi*^*ts*^ in their PRCs and shifted to the restrictive temperature did not show axonal degeneration during prolonged light exposure (Fig. 5E), even after 13 days of constant light exposure, 13dLL (Fig. 5E, F). This suggests that blocking neurotransmitter release protects against axonal degeneration.

**Figure 5:**
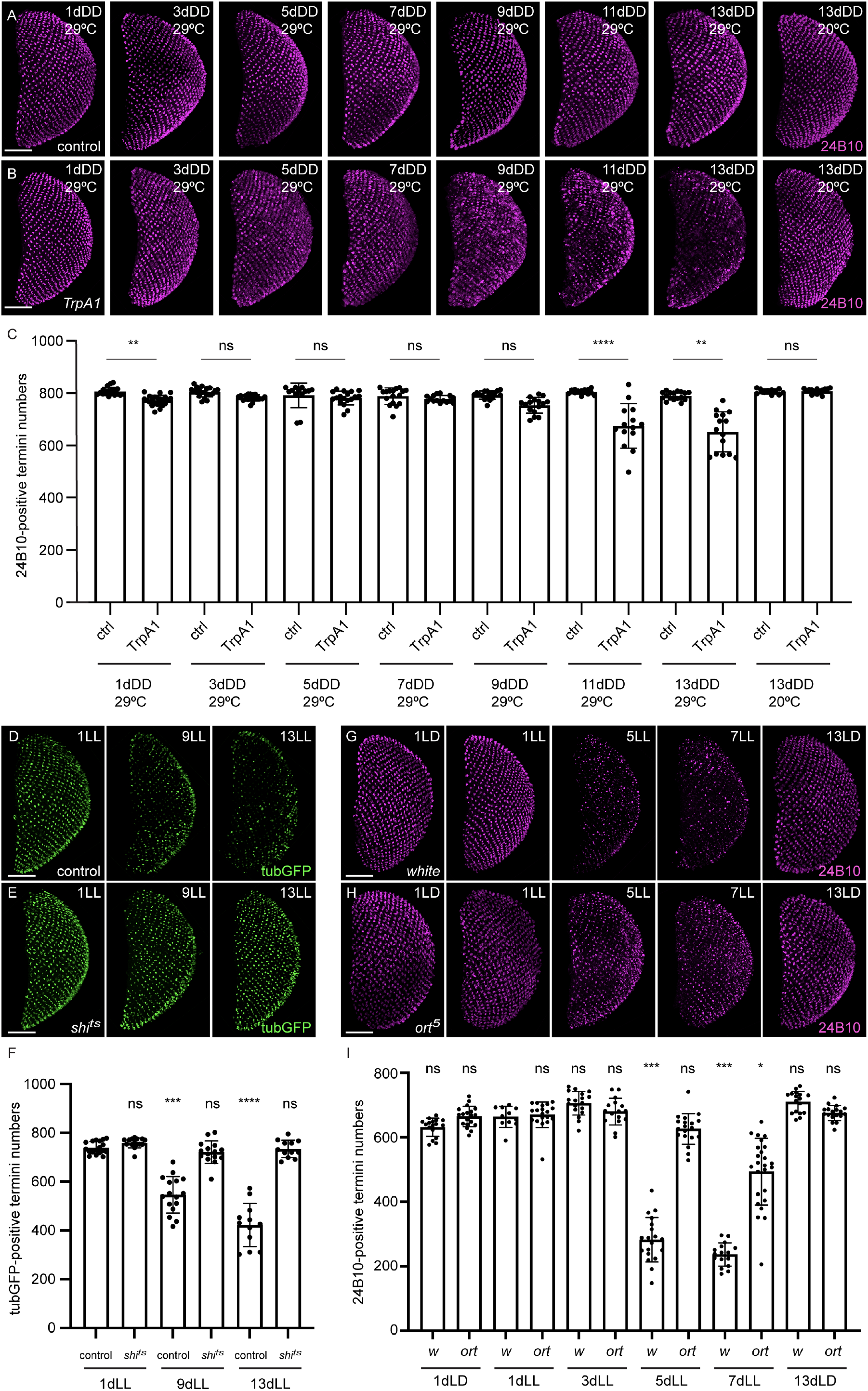
Neurotransmission is involved in the axonal degeneration process. **A, B**. Axonal termini of R7 in the medulla of flies expressing *UAS-TrpA1* or control flies kept in complete darkness (DD) for various periods of time, either at 29ºC (test condition) or at 20 ºC (control). Magenta represents termini stained with 24B10. Scale bar: 50 μm. **C**. Graph of 24B10-positive R7 termini counts in the medulla of *UAS-TrpA1*-expressing or control flies kept in complete darkness (DD) for various periods of time, either at 29ºC (test condition) or at 20 ºC (control). **D, E**. Axonal termini of R7 in the medulla of flies expressing *UAS-shi*^*ts*^ in (E) and control flies in (D) exposed for 1, 9 or 13 days to constant light (LL). Green represents termini expressing *UAS-tubulinGFP* driven by *GMR-gal4*. Scale bar: 50 μm. **F**. Graph of tubulinGFP-positive R7 termini counts in the medulla of flies expressing *UAS-shi*^*ts*^ or control flies exposed for 1, 9 or 13 days to constant light (LL). **G, H**. Axonal termini of R7 in the medulla of *white* flies in (G) or flies mutant for *ort* in (H) exposed for various times to constant light (LL) or to control light conditions for 1 or 13 days (LD). Magenta represents termini stained with 24B10. Scale bar: 50 μm. **I**. Graph of 24B10-positive R7 termini counts in the medulla of control *white* flies or *ort* mutant flies exposed for 1, 3, 5 or 7 days to constant light (LL) or kept for 1 or 13 days in LD.

These results implied that R7 axonal degeneration required the involvement of postsynaptic neurons. To directly test this and to rule out potential additional effects of hindering endocytosis by Shi^ts^, we blocked the postsynaptic response to PRC activation, leaving PRC activity intact. Fly photoreceptors are histaminergic and the Ort histamine chloride channel promotes hyperpolarization of R7 postsynaptic neurons in the medulla (Gao et al., 2008; Gisselmann et al., 2002; Liu and Wilson, 2013; Pantazis et al., 2008; Schnaitmann et al., 2018). However, we did not succeed in establishing a line containing an *ort* mutant (Gengs et al., 2002) in the *GMR white RNAi* background. As an alternative, we therefore included the *ort*^*5*^ allele in *white* mutant flies. Exposure of adult *white* flies to constant light gave rise to a very strong degeneration after only 5 days of exposure, as detected by 24B10 staining (Fig. 5G, I, compare to Fig. 2J, K). With this experiment, we thus confirmed previous results obtained in the retina, showing that pigments protect photoreceptor cell bodies from excessive exposure to light (Bulgakova et al., 2010; Ferreiro et al., 2017; Lee and Montell, 2004). The presence of *ort*^*5*^ protected flies against axonal degeneration, in fact decreasing the loss of axons after 5 and 7 days of constant light exposure (Fig. 5H, I). Together with the *shi*^*ts*^ data, this result showed that neurotransmission to postsynaptic partners is required to initiate R7 axon degeneration. It further suggested that prolonged hyperpolarization of the postsynaptic neurons represents an important signal for the initiation of axonal degeneration in R7.

### Synapse loss precedes axonal degeneration

Synaptic dysfunction is considered as an early event and as the major determinant of ND, including AD and PD (Bellucci et al., 2016; Colom-Cadena et al., 2020; Compta and Revesz, 2021). We thus asked whether synaptic function was also affected in our model. For this, we took advantage of the activity-dependent syb:GRASP system, which allows for retrospective labelling of synapses based on their activity (Macpherson et al., 2015). We expressed the *syb-spGFP1–10* construct in the *yellow* ommatidia subset of R7s using *Rh4lexA* as a driver and *CD4-spGFP11* in Dm8, a synaptic partner of R7 in the medulla, using *ortGal4* (Gao et al., 2008; Schnaitmann et al., 2018). Only the combination of the two fragments across the plasma membrane of active synapses yields a detectable GFP signal (Macpherson et al., 2015). These flies were then subjected to a control LD treatment (1 day and 13 days LD) or put in LL for different periods of time (1, 5, 7, 9, 11 or 13 days). At every timepoint, 24B10-positive axonal termini were counted (Fig. 6E) and compared to numbers of GFP-positive termini, which represent active synapses between the *yellow* R7 PRCs and Dm8 (Fig. 6F). We counted a total number of approximately 750 R7 axonal termini labeled with 24B10 per medulla (Fig. 6E) as well as ca. 450 GRASP-positive termini (Fig. 6F) in control conditions (1dLD), corresponding to the approximate expected fraction of *yellow* ommatidia (70% of total R7) (Franceschini et al., 1981). These numbers remained constant over time in an LD cycle (Fig. 6E, F). In this genotype, 24B10-positive termini were unchanged at 7dLL (Fig. 6A, C, E and Suppl. Fig. 4A, D). Axonal degeneration in this genotype started at 9dLL (Fig. 6D, E and Suppl. Fig. 4E) and proceeded progressively in the following days (Fig. 6E). In contrast, GRASP counts were already clearly reduced at 7 days in LL (drop from ca. 450 to ca. 300 GFP-positive termini) (Fig. 6B’, C’, F), thus before axonal degeneration was detectable.

**Figure 6.**
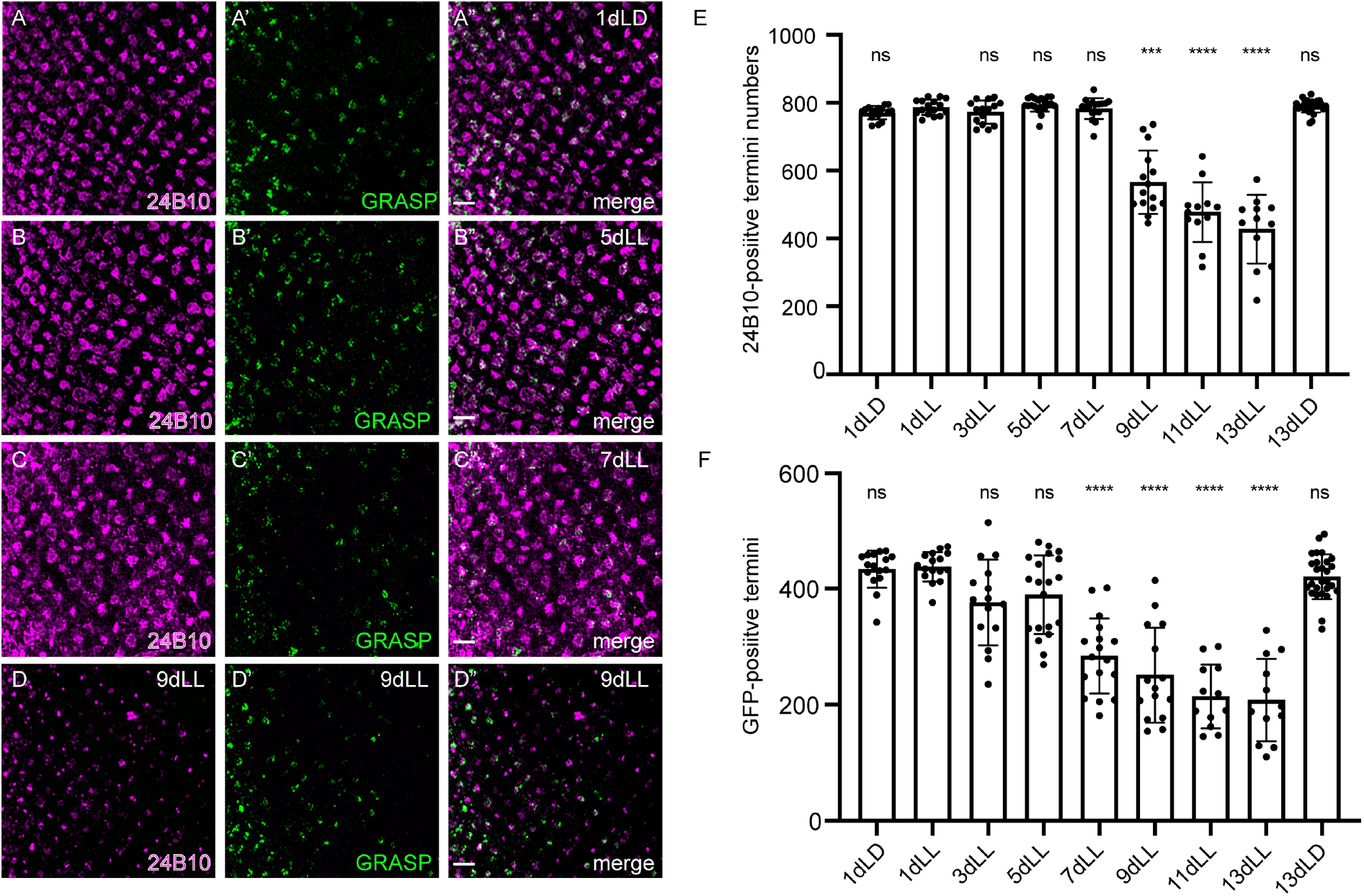
Synaptic dysfunction between R7 and Dm8 precedes axonal degeneration. **A-D”**. Close-ups on the axonal termini of R7 in the medulla of flies exposed for various time to constant light (5 days in B-B”; 7 days in C-C”; 9 days in D-D”) or to control condition for 1 day in a 12 hours of light/12 hour of darkness (LD) condition (A-A”). (A-D) show the termini stained with chaoptin (24B10) in magenta. (A’-D’) depict sybGRASP-positive contacts between R7 and Dm8 (green). Last column (A”-D”) shows a merge of the close-ups for 24B10 and GFP. Scale bar: 10 μm. **E**. 24B10-positive R7 termini in the medulla of flies exposed for various time to constant light (LL) or control conditions (LD). **F**. GRASP-positive R7 termini numbers in the same medullas as in E.

Taken together, these data indicate that activity-dependent synaptic transmission between R7 and a medullar postsynaptic partner is lost before axonal termini numbers start to decrease, suggesting that loss of synaptic connection precedes axonal degeneration.

Thus, the model system of sporadic axonal degeneration that we developed shares important characteristics with the reported progression of neurodegeneration in human ND.

## Discussion

In this work, we established a model of sporadic initiation of axonal degeneration. We describe here the amenability of using R7 photoreceptors axonal termini in the medulla as a readout for sporadic axonal degeneration. For this, we developed tools for the quantitative analysis of R7 axon loss in the medulla. With this system, we defined that the stage of initiation of axonal degeneration is reached individually for each neuron. Furthermore, we showed that synapses between R7 and a postsynaptic partner are dysfunctional before axonal degeneration starts. We thus produced a model system, in which we recapitulate important early features of human neurodegenerative diseases. In addition, we provided evidence for a postsynaptic signal regulating stability of the presynaptic axon. Thus, the model we established can now be used to define the early changes that happen within neurons at the time point in which their resilience capacity is overcome.

Various factors can affect axon vulnerability, including cell senescence, metabolic changes, neuroinflammation and exposure to chemicals, including some utilized in cancer therapy (Adalbert and Coleman, 2013; Coleman, 2005; Figley and DiAntonio, 2020; Neukomm and Freeman, 2014). In this model, we induced axonal degeneration by stimulating neurons for prolonged periods of time. Photoreceptors are resistant to prolonged stimulation, thanks to a series of mechanisms acting on a short time scale to maintain their response and output within a dynamic range and potentially to maintain circuit homeostasis on a longer time scale (Juusola and Hardie, 2001; Stavenga, 1995; Sugie et al., 2015). Thus, they represent a good system to investigate how this resilience is no longer sufficient to guarantee axon maintenance. Further, imbalanced activity represents a potential trigger of neurodegeneration (Busche and Konnerth, 2016; Palop and Mucke, 2016; Palop et al., 2007; Sosulina et al., 2021). Neuronal hyperactivity has been detected in patients with mild cognitive impairment (MCI), a prodromal stage of AD, and in carriers of the APOE4 allele, the most important genetic risk factor for late-onset sporadic AD, as well as in many transgenic AD mice (Hector and Brouillette, 2020). Furthermore, activity imbalance at the circuit level is reported in Alzheimer’s disease and in AD model systems and is considered a potential trigger of neurodegeneration (Busche and Konnerth, 2016; Palop and Mucke, 2016; Palop et al., 2007; Sosulina et al., 2021). In addition, the exacerbated action of excitatory neurotransmitters, primarily glutamate, is reported to contribute to the neurotoxicity observed in major neurodegenerative diseases (Armada-Moreira et al., 2020; Salińska et al., 2006). Our present data suggest that PRC axon degeneration is not induced by excitotoxicity, since it can be rescued to a large extent by blocking synaptic vesicle release or even by blocking the response of postsynaptic neurons. Thus, we propose that an unbalanced level of activity within the local microcircuit at the PRC output synapses is at the core of the transition towards initiation of axonal degeneration in this system. The signals that trigger the onset of degeneration in this context remain to be identified and could be counterbalancing the pathways that initially maintain local homeostasis (Sugie et al., 2015).

Axonal degeneration induced in constant light conditions does not affect a particular region of the medulla. This stochastic start of degeneration is preceded by the swelling of R7 axonal termini, which takes place throughout the medulla. This change appears similar to that reported in degenerating axons in the central nervous system of rodents (Ertürk et al., 2007). The swelling is reversed by placing the animals back in the dark for prolonged periods of time. In contrast to the swelling, which was shared by all axons, only individual axons lost their tubulin and membrane markers and degenerated. At 7dLL, the first axons started degenerating, but putting 7dLL animals back to DD blocked further axonal loss, indicating that the timing of trigger, rather than the timing of execution of an axonal degeneration program is different among PRCs. This suggests that the resilience capacity of a single axon is different from that of its neighbours and it will be crucial to understand the mechanisms underlying this phenomenon.

We observed that synapses between R7 and one of its postsynaptic partners in the medulla lose their integrity before axons degenerate. These results raise the question whether synaptic detachment could be causative of axonal degeneration or whether synaptic maintenance mechanisms were lost before synapses detached. A broad line of evidence suggests that alterations in synaptic adhesion play key roles in the disruption of neuronal networks in neurodegenerative diseases (Chapman, 2014; Kilinc, 2018; Leshchyns’ka and Sytnyk, 2016). Synaptic maintenance needs to cope with the metabolic demand of neurotransmission, as well as with elevated rates of protein turnover and a high membrane exchange that requires efficient delivery and constant supply of newly synthesized proteins (Guedes-Dias and Holzbaur, 2019; Harris et al., 2012). Therefore, removal of damaged proteins and organelles from synaptic sites is essential to sustain synaptic function (Andres-Alonso et al., 2021). Abnormalities in function, trafficking or signaling of mitochondria, lysosomes and endoplasmic reticulum contribute to development of neurodegenerative diseases (Cabral-Miranda and Hetz, 2018; Kerr et al., 2017; Lie and Nixon, 2019; Öztürk et al., 2020). It will thus be of high interest to clarify whether axonal transport deficiencies, ER stress, mitochondrial and/or lysosomal dysfunction precede synaptic detachment in this model.

We found no signs of apoptosis in R7 up to 13 days of exposure to constant light, when axonal degeneration was very advanced. Thus, in this model, axonal degeneration precedes cell death of several days, reproducing the sequence of events observed in human neurodegenerative diseases (Coleman, 2005). Interestingly, we observed axonal swellings and microtubule disassembly upon light or TrpA1 treatment, both features often preceding cell body loss in the ‘dying back pathology’ described in many neurodegenerative diseases (Wang et al., 2012). Increasing evidence shows that injury-induced degeneration (Wallerian degeneration) shares molecular features with neurodegenerative diseases’ ‘dying back’ axonal degeneration: the local loss of NMNAT2 activates either SARM1 Wallerian degeneration or triggers SARM1-dependent ‘dying back’ (Coleman and Höke, 2020; Figley and DiAntonio, 2020; Yaron and Schuldiner, 2016). For this reason, it would be crucial to address the involvement of NMNAT2, SARM1 and other players of this pathway in our model.

Taken together, we developed a quantitative stochastic model of axonal degeneration. Our initial characterization indicates that it reproduces important traits of human neurodegenerative diseases, including the interruption of neuronal resilience to repetitive activation and the limitation of the initial defects to the axons, without resulting in immediate cell death. Our results further point to a role of circuit imbalance towards the initiation of axonal degeneration.

## Acknowledgments

This work was supported by the DZNE core funding (GT) and by grants from the Ministry of Education, Culture, Sports, Science and Technology of Japan (#18K14835, #18J00367 and #21K15619 to YN, #17H04983, #19K22592 and #21H02837 to AS), and Takeda science foundation life science research grant to AS. We thank R. Kerpen, P. Tran and O. Sharma for technical support. We are grateful to the Tavosanis lab and to F. Bradke, D. Di Monte and E. Mandelkow for helpful discussions and comments on the manuscript. We gratefully acknowledge Profs. T. Lee, M. Pankratz and H. Tanimoto, BDSC (NIH P40OD018537) and VDRC for providing flies. We thank the Virtual fly brain for the 3D brain stacks (Fig. 1B, C) (https://virtualflybrain.org).

## Supplemental Tables

**Supplemental table 1.**
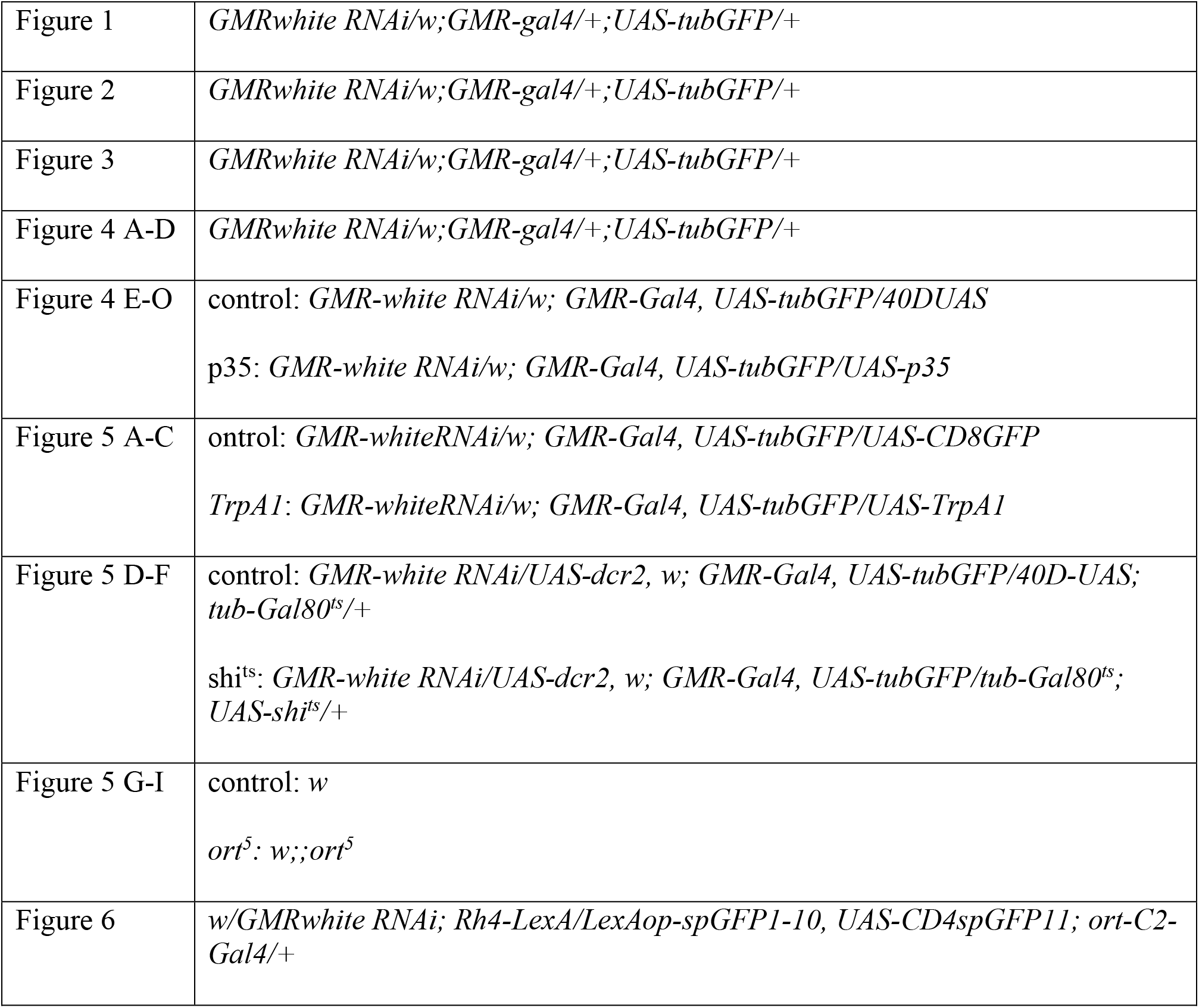
Genotypes

**Supplemental table 2.**
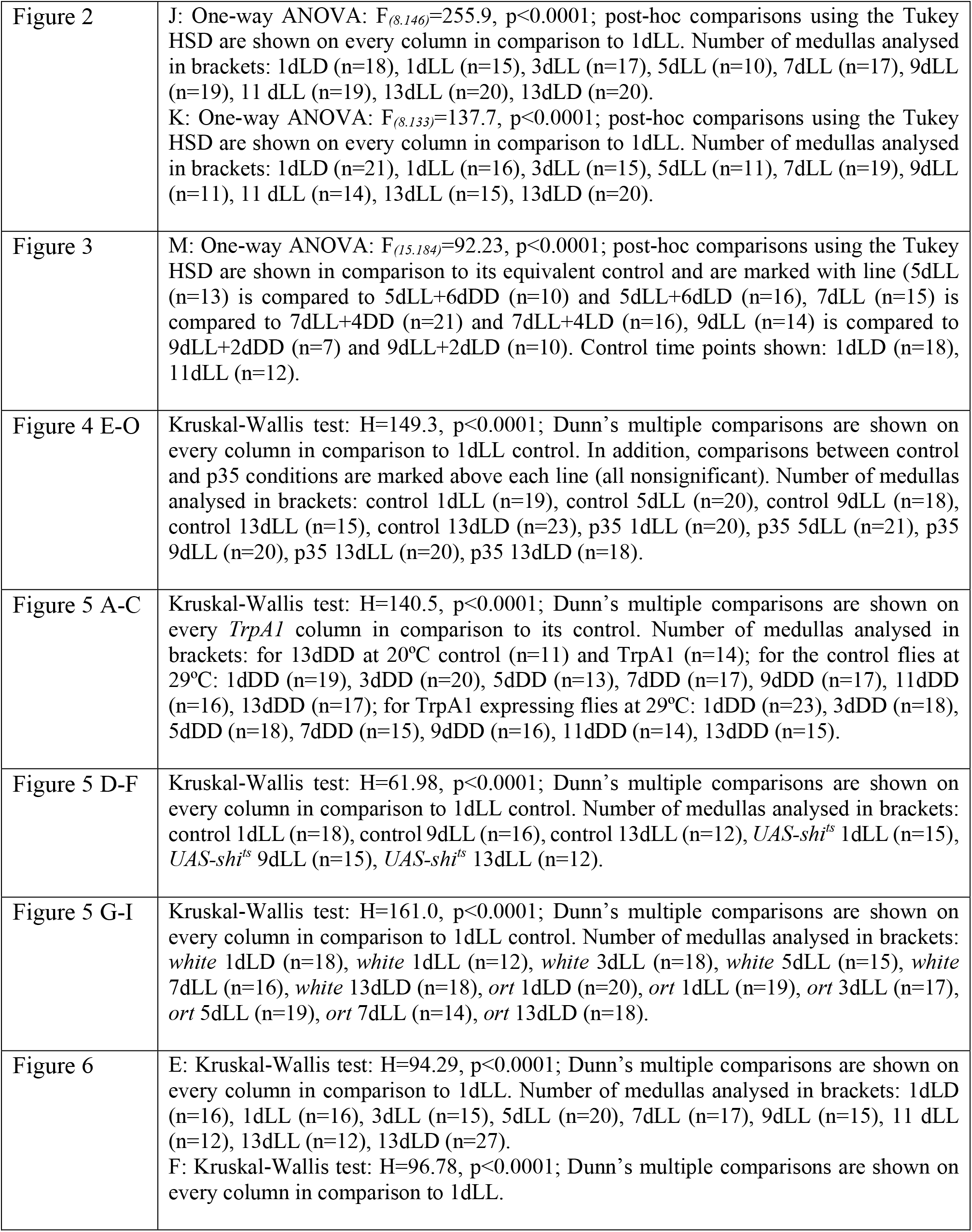
Statistics

## Supplemental figures

**Supplemental Figure 1.**
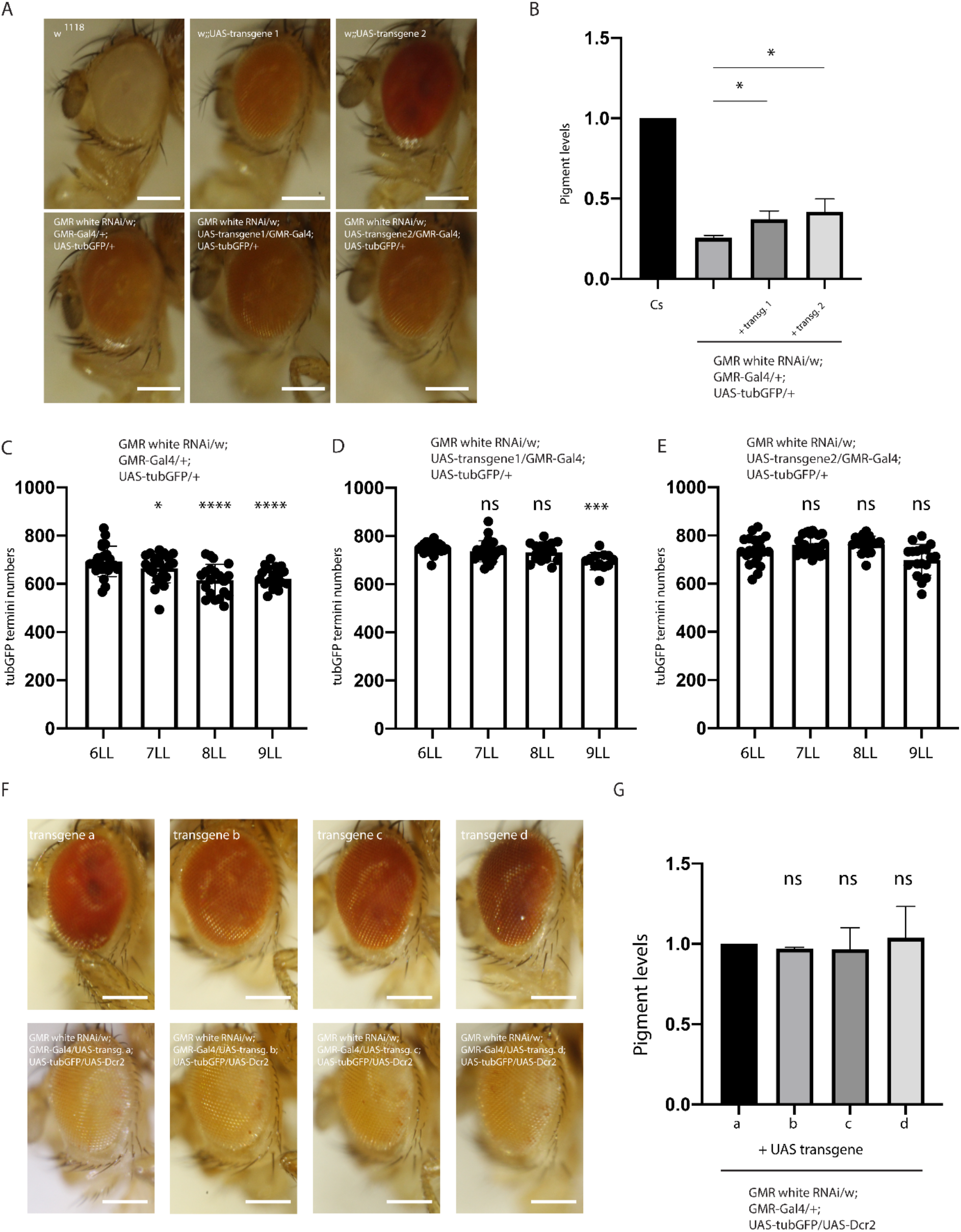
Eye pigment content influences the time of axon degeneration onset. **A, B**. The presence of additional transgenes in the *GMR white RNAi/w; GMRgal4 UAS-GFP* background changes eye color to a certain extent. (A) Eye of a *w*^*1118*^ fly (top left) or of two different lines including *white*^*+*^-marked UAS transgenes (transgene 1 and 2) in a *w*^*1118*^ background, leading to different eye colors (top right panels). The bottom panels show eyes of *GMR white RNAi/w;GMR-Gal4/+;UAS-tubGFP/+* (bottom left); of *GMR white RNAi/w; UAS-transgene1/GMR-Gal4;UAS-tubGFP/+* (bottom center) or *GMR white RNAi/w; UAS-transgene2/GMR-Gal4;UAS-tubGFP/+* (bottom right). Scale bar: 250 μm. (B) Brown pigments content in eyes of 7-10 days old female flies *GMR white RNAi/w;GMR-Gal4/+;UAS-tubGFP/+, GMR white RNAi/w; UAS-transgene1/GMR-Gal4;UAS-tubGFP/+* or *GMR white RNAi/w; UAS-transgene2/GMR-Gal4;UAS-tubGFP/+* in comparison to wild-type *Canton S* levels. Bars represent mean ±SD of 3 experiments. Pigment levels in *Canton S* were arbitrary set to 1.0. One-way ANOVA: F_*(3*.*8)*_=143.1, p<0.0001; post-hoc comparisons using the Tukey HSD are shown above the columns. **C-E**. A darker eye color is associated with a slower axonal degeneration. TubulinGFP-positive R7 termini counts in the medulla of *GMRwhite RNAi/w;GMR-gal4/+;UAS-tubGFP* flies (C) *GMR white RNAi/w; UAS-transgene1/GMR-Gal4;UAS-tubGFP/+* (D) as well as *GMR white RNAi/w; UAS-transgene2/GMR-Gal4;UAS-tubGFP/+* (E) exposed for various time to constant light (LL). (C) One-way ANOVA: F_*(3*.*80)*_=11.31, p<0.0001; post-hoc comparisons using the Tukey HSD are shown on every column in comparison to 6dLL. Number of medullas analysed in brackets: 6dLL (n=23), 7dLL (n=23), 8dLL (n=20), 9dLL (n=18). (D) One-way ANOVA: F_*(3*.*70)*_=5.682, p=0.0015; post-hoc comparisons using the Tukey HSD are shown on every column in comparison to 6dLL. Number of medullas analysed in brackets: 6dLL (n=22), 7dLL (n=23), 8dLL (n=17), 9dLL (n=12). (E) One-way ANOVA: F_*(3*.*78)*_=10.80, p<0.0001; post-hoc comparisons using the Tukey HSD are shown on every column in comparison to 6dLL. Number of medullas analysed in brackets: 6dLL (n=20), 7dLL (n=25), 8dLL (n=19), 9dLL (n=18). **F, G**. The presence of a *UAS-Dcr2* transgene leads to a homogeneous eye color independently of the addition of *w*^*+*^*-*expressing transgenes. F: The upper row shows from left to right: *w;UAS-transgene a/CyO;UAS-Dcr2/TM6B* or *w;UAS-transgene b/CyO;UAS-Dcr2/TM6B* or *w;UAS-transgene c/CyO;UAS-Dcr2/TM6B* or *w;UAS-transgene d/CyO;UAS-Dcr2/TM6B*. The bottom lane illustrates, from left to right, adult eyes of: *GMR white RNAi/w;GMR-Gal4/UAS-transgene a;UAS-tubGFP/UAS-Dcr2* or *GMR white RNAi/w;GMR-Gal4/UAS-Dcr2;UAS-tubGFP/UAS-transgene b* or *GMR white RNAi/w;GMR-Gal4/UAS-transgene c;UAS-tubGFP/UAS-Dcr2* as well as *GMR white RNAi/w;GMR-Gal4/ UAS-transgene d;UAS-tubGFP/UAS-Dcr2*. Scale bar: 250 μm. G. Brown pigments content in eyes of 7-10 days old female flies illustrated in bottom panels in F. Bars represent mean±SD of 3 experiments. Pigment levels in the *GMR white RNAi/w;GMR-Gal4/UAS-transgene a;UAS-tubGFP/UAS-Dcr2* flies were arbitrary set to 1.0. Eye pigment contents of *GMR white RNAi/w;GMR-Gal4/UAS-transgene b;UAS-tubGFP/UAS-Dcr2* or *GMR white RNAi/w;GMR-Gal4/UAS-transgene c;UAS-tubGFP/UAS-Dcr2* as well as *GMR white RNAi/w;GMR-Gal4/ UAS-transgene d;UAS-tubGFP/UAS-Dcr2* are plotted. One-way ANOVA: F_*(3*.*8)*_=0.23, p=0.8681; post-hoc comparisons using the Tukey HSD are shown in comparison to the first column.

**Supplemental Figure 2.**
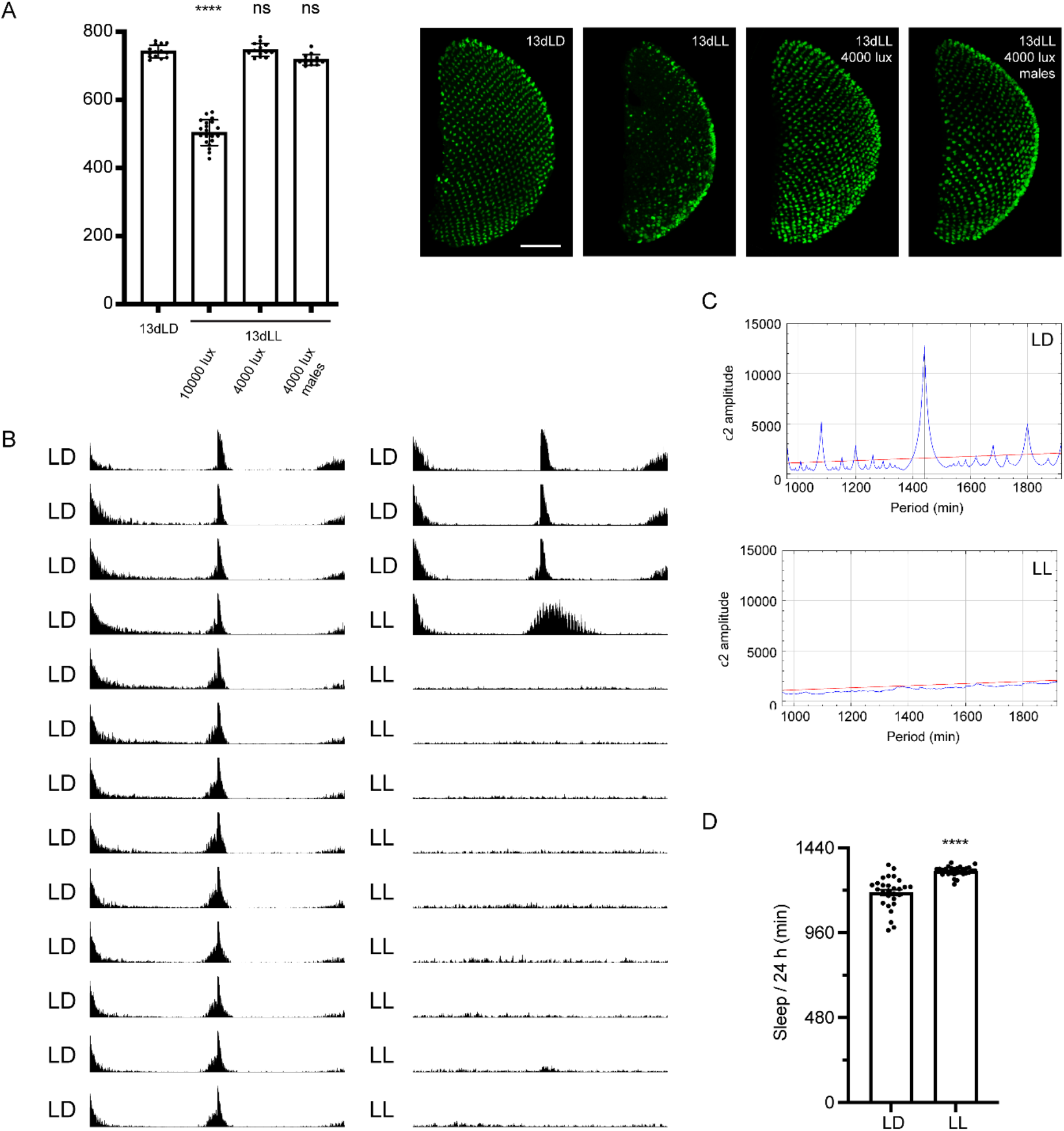
Disruption of circadian rhythm or sleep is not the cause of axonal degeneration. **A**. TubulinGFP-positive R7 termini counts in the medulla of *GMRwhite RNAi/w;GMR-gal4/+;UAS-tubGFP* flies exposed to 13 days LD (first column) or 13 days in constant light of 10000 lux (second column) or 4000 lux (third column), or males *GMR white RNAi/Y;GMRgal4 UAS-tubGFP*/+ exposed to 13 days in constant light of 4000 lux (fourth column). One-way ANOVA: F_*(4*.*68)*_=282.3, p<0.0001; post-hoc comparisons using the Tukey HSD are shown on every column in comparison to 13dLD. Number of medullas analysed in brackets: 13dLD (n=14), 13dLL 10000 lux (n=17), 13dLL 4000 lux (n=10), 13dLL 4000 lux males (n=11). An exemplary picture of *tubGFP*-labelled axonal termini of R7 in the medulla is shown on the right for each condition. Scale bar: 50 μm. **B**. Actogram showing the average locomotor activity of *GMR white RNAi/Y;GMRgal4 UAS-tubGFP*/+ flies in light 12h/ dark 12h cycle (LD, left) and constant light (LL, right). Flies were entrained to a 12-h:12-h LD cycle (light: 4000 lux) at 25°C for 3 days (upper 3 actograms). Subsequently, test flies were subjected to constant light (LL) conditions for 10 days and control flies to LD cycles for 10 days. **C**. χ^2^ periodogram records of LD (upper) or LL (lower) conditions in *GMR white RNAi/Y;GMRgal4 UAS-tubGFP*/+ flies. The blue line shows a period of 1440 min (24h) for flies in LD while no periodicity was detected in LL flies. The red line indicates a significant level of *p* = 0.05. **D**. Total amount of sleep of *GMR white RNAi/Y;GMRgal4 UAS-tubGFP*/+ flies exposed to LD (n=28) and LL (n=28) at an intensity of 4000 lux during 24 hours (Mann-Whitney U-test: P<0.0001). Sleep was defined as an absence of movement for more than 5 minutes.

**Supplemental Figure 3.**
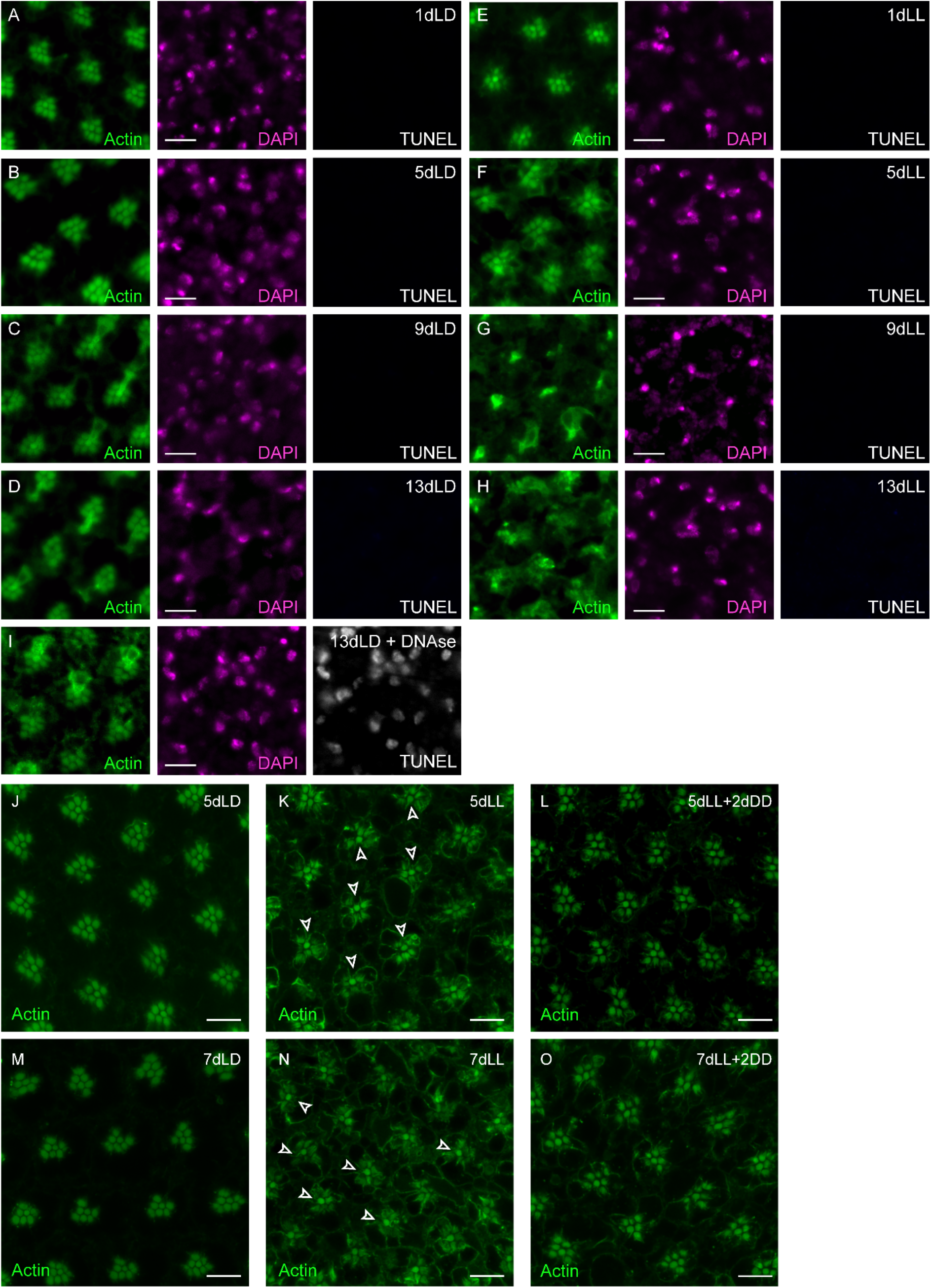
Apoptosis is not associated to axonal degeneration and rhabdomere remodeling upon light exposure is reversible. **A-I**. Optical cross-sections through the retina of *GMR white RNAi/w;GMR-Gal4/+;UAS-tubGFP/+* adult flies exposed to control (LD) or constant (LL) light conditions for 1, 5, 9 or 13 days. Eyes were stained with FITC-Phalloidin to highlight the actin-enriched rhabdomeres (green) and DAPI for the nuclei (magenta). TUNEL labelling showed no signal except in the positive control, in which 13dLD eyes were submitted to a DNAse treatment before TUNEL detection (I). Scale bar: 10 μm. **J-O**. Optical cross-sections through the retina of *GMR white RNAi/w;GMR-Gal4/+;UAS-tubGFP/+* adult flies exposed to control (LD) or constant (LL) light conditions for 5 (J, K) or 7 days (M, N) or exposed to 5 or 7 days of constant light following 2 days of constant darkness (L, O). Eyes were stained with FITC-Phalloidin to highlight the actin-enriched rhabdomeres (green). Arrowheads point to R7 in K and N. Scale bar: 10 μm.

**Supplemental Fig. 4.**
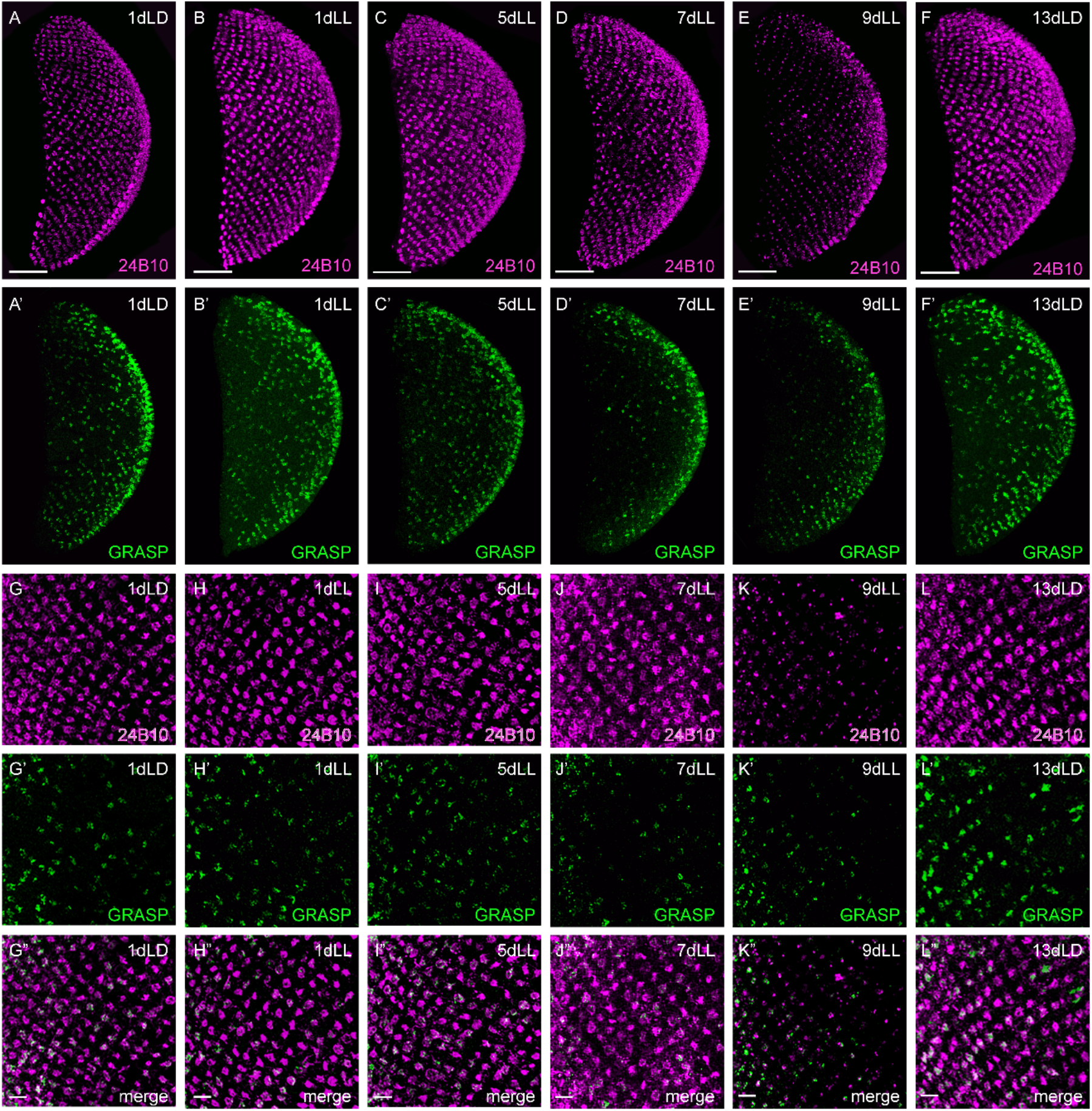
Synaptic dysfunction between R7 and Dm8 precedes axonal degeneration. **A-L”**. Axonal termini of R7 in the medulla of *w/GMRwhite RNAi; Rh4-LexA/LexAop-spGFP1-10, UAS-CD4spGFP11; Ort-C2-Gal4/+* flies exposed for various time to constant light (1 day in B, B’ and a close-up in H-H”; 5 days in C, C’ and a close-up in I-I”; 7 days in D, D’ and J-J”; 9 days in E, E’ and the close-up in K-K”) or to control condition for 1 day in a 12 hours of light/12 hour of darkness (LD) condition (A, A’ and G-G” for a close-up) as well as 13 days in LD in F, F’ and L-L”. **A-L** Axon termini labeled with chaoptin (24B10) in magenta. **A’-L’** depict GFP-positive termini in which a GRASP event took place between R7 and Dm8 (green). **G”-L”** shows a merge of the close-ups for 24B10 and GFP. Scale bar for the whole medullas: 50 mm (A-F’) and for the close-ups: 10 μm (G-L”).

